# Lipidomic Profiling of L1CAM+ Extracellular Vesicles in Parkinson’s Disease

**DOI:** 10.64898/2026.11.21.768278

**Authors:** Mohammed Adnan Abdulsalam Abdulsalam, Cemil Can Eylem, Izzet Avci, Hare Yazgi, Merve Akkulak, Cansu Beyret, Hasret Gonul, Hakan Eskizengin, Kerem Atalar, Mehmet Atakay, Emirhan Nemutlu, Bekir Salih, Mehmet Gokhan Caglayan, Rezzak Yilmaz, Burak Derkus

## Abstract

Parkinson’s disease (PD) lacks minimally invasive biomarkers that reflect neuron-derived molecular alterations. Here, we performed a pilot study for untargeted Liquid Chromatography-Mass Spectrometry (LC-MS)-based lipidomics on serum total extracellular vesicles (tEVs) and L1CAM-enriched EVs (L1CAM+EVs) in patients with PD and healthy controls (n=10 per group; N=40). Partial least squares-discriminant analysis (PLS-DA) separated PD from control samples in both fractions L1CAM+EVs: R^2^Y = 0.998, Q^2^ = 0.879; tEVs: R^2^Y = 0.998, Q^2^ = 0.876), and distinguished L1CAM+EVs from tEVs within PD samples (R^2^Y = 0.999, Q^2^ = 0.925). In PD tEVs, triacylglycerols, fatty acids, phosphatidylcholines, sphingomyelins and phosphatidylserines were increased, and pathway analysis indicated enhanced conversion of phosphatidylcholine (PC) and phosphatidylethanolamine (PE) to phosphatidylserine (PS). In contrast, PD L1CAM+EVs showed increased DG, LPA and PS with reduced LPC; pathway reconstruction indicated enhanced PE-PC-DG and LPC-PC flux with reduced de novo PC and SM synthesis. Within PD, L1CAM+EVs were depleted of multiple sphingomyelin species (e.g., SM 41:2;O3, SM 35:2;O3, SM 41:3;O3) and glycerophospholipids relative to tEVs, whereas SPB 18:0;O2 and FA 16:4 were relatively enriched. These findings indicate that L1CAM-enriched EVs carry a PD-associated lipid signature that is qualitatively distinct from that of controls, supporting their evaluation as a complementary source of neuron-related lipid biomarkers.

## 1. INTRODUCTION

Parkinson’s disease (PD) is the second most common neurodegenerative disorder worldwide, defined by the progressive loss of dopaminergic neurons in the substantia nigra pars compacta and the accumulation of misfolded α-synuclein in Lewy bodies and Lewy neurites (**1,2**). Diagnosis still relies on clinical examination, and biomarkers that reflect ongoing neuronal pathology remain an unmet need (**3**). On that account, α-synuclein seed amplification assays (SAA) have recently transformed the field: they detect pathological α-synuclein with high sensitivity and specificity, including in prodromal stages, and now underpin proposed biological definitions of the disease (**4–7**). However, SAA provides a largely qualitative readout, is performed mainly on cerebrospinal fluid, and offers limited information on the metabolic state of affected neurons or on disease progression. Complementary, quantitative and blood-accessible measures of neuronal biology are therefore still needed.

Lipid dysregulation is increasingly recognized as a central component of PD pathogenesis (**8,9**). Neuronal function depends on tightly regulated lipid homeostasis governing synaptic vesicle dynamics, membrane trafficking, mitochondrial integrity and lysosomal function (**10**). Alterations in sphingolipids, glycerophospholipids and fatty acids have been reported in PD brain tissue and in blood (**11,12**). α-Synuclein is itself a lipid-binding protein: its conformation, membrane affinity and aggregation propensity are strongly modulated by the surrounding lipid environment, particularly by anionic phospholipids and membrane curvature (**13,14**). Genetic evidence points in the same direction. Variants in GBA1, encoding glucocerebrosidase, are among the strongest genetic risk factors for PD and directly link sphingolipid metabolism to disease susceptibility (**15,16**). Altered sphingomyelin-ceramide balance and ceramide acyl-chain composition have also been described in PD brain (**17**). In addition, mitochondrial and oxidative stress can drive the remodeling of phospholipid and fatty acid species (**18**).

Extracellular vesicles (EVs) provide a means of sampling these processes. Lipids are fundamental determinants of EV biology, shaping membrane rigidity and curvature, vesicle biogenesis, cargo sorting, circulatory stability and uptake by recipient cells (**19–21**). Sphingolipid metabolism is directly coupled to vesicle formation: ceramide generated by neutral sphingomyelinase promotes ESCRT-independent budding of intraluminal vesicles (**22**). Sphingomyelin and phosphatidylcholine/phosphatidylethanolamine species, in turn, define the order and architecture of the vesicle bilayer (**23**). Importantly, EVs from patients with Lewy body disorders show altered ceramide metabolism linked to EV-mediated spread of α-synuclein (**24**). Changes in EV lipid composition may therefore reflect not only the metabolic state of the parental cell, but also processes involved in pathological propagation.

Interpreting blood-derived EV or lipid signatures, however, is challenging, because circulating lipids and vesicles originate from many organs and cell types (**25**). Neuron-specific alterations may thus be diluted within the peripheral lipidome. To address this limitation, immunocapture of EVs carrying the neural cell adhesion molecule L1CAM has been widely used to enrich putatively neuron-derived EVs from plasma or serum (**26,27**). In PD, L1CAM-enriched EVs contain elevated α-synuclein, which may help distinguish PD from atypical parkinsonism, and identify individuals at risk of developing PD (**28–30**). Similar approaches have been applied in Alzheimer’s disease and other neurological disorders (**31,32**).

However, while the protein and RNA cargo of L1CAM-enriched EVs has been studied extensively, their lipid composition in PD remains poorly characterized. It is also unknown whether L1CAM enrichment captures disease-associated lipid alterations that are less evident in total circulating EVs. Here, we performed untargeted lipidomics on serum total EVs (tEVs) and L1CAM-enriched EVs (L1CAM+EVs) from patients with PD and matched healthy controls. Combining multivariate modelling, differential abundance and lipid-class analyses with reaction-network and pathway reconstruction, we aimed to (i) define PD-associated lipid alterations in each EV fraction, (ii) compare the lipid composition of L1CAM-EVs and tEVs within PD samples, and (iii) assess whether L1CAM enrichment improves the discrimination of PD from controls relative to tEVs.

## 2. MATERIALS and METHODS

### 2.1. Study Design

The study included individuals recruited as part of the ANkara PArkinson’s Disease Registry (ANPAR) cohort Patients with PD (n=20) and age-matched healthy control individuals (n=20) were randomly allocated to tEV or L1CAM+EV isolation (n=10 per group; N=40). Ethical approval was obtained from the local ethics committee (approval no. i04-327-25), and the study was conducted in accordance with the ethical principles of the Declaration of Helsinki. Demographic and clinical characteristics of the study participants are presented in **Table 1**. Total serum EVs (tEVs) were first isolated by sequential differential centrifugation followed by ultracentrifugation (UC). Subsequently, an L1CAM-enriched EV fraction was generated from the purified tEV pool using biotinylated anti-L1CAM antibody-based immunocapture. This two-step workflow, consisting of UC-derived tEV isolation followed by L1CAM-positive EV enrichment, was conceptually adapted from the method reported by Mustapic et al. (**29**), with several optimizations including pooled tEV input, modified incubation conditions, and mass spectrometry-compatible buffer adjustments. The study was structured in two analytical stages. First, the lipidomic composition of L1CAM-enriched EVs was directly compared with matched tEV fractions to determine whether neuronal-associated enrichment provides a distinct lipid signature relative to the total circulating EV pool. Second, disease-associated comparisons were performed between Parkinson’s disease-derived EVs and control-derived EVs within each fraction type (tEV and L1CAM-enriched EVs).

**Table 1.** Demographic and Clinical Characteristics of the Study Groups.

| **Variable** | **Parkinson’s Disease** | **Healthy Control** | **p** |
|---|---|---|---|
| **tEV Group** |  |  |  |
| **n** | 10 | 10 | - |
| **Age, mean ± SD** | 61.4 ± 8.5 | 58.8 ± 11.0 | 0.256 |
| **Male sex n, (%),** | 5 (50) | 5 (50) | 1.000 |
| **BMI, mean ± SD** | 26.1 ± 3.4 | - | - |
| **Disease duration (years), median? (range)** | 5.5 (3.5 - 7.8) | - | - |
| **Hoehn & Yahr stage** | 2.0 (2.0 - 2.8) | - | - |
| **MDS-UPDRS III** | 41.5 (23.5 - 55.2) | - | - |
| **LEDD (mg/day)** | 655.8 (387.5 - 850.0) | - | - |
| **L1CAM+ EV Group** |  |  |  |
| **n** | 10 | 10 | - |
| **Age, mean ± SD** | 61.1 ± 3.3 | 60.3 ± 6.0 | 0.475 |
| **Male Ssex n, (%)** | 5 (50) | 7 (70%) | 0.630 |
| **BMI, mean ± SD** | 31.0 ± 5.7 | - | - |
| **Disease duration (years)median? (range)** | 3.0 (0.2 - 5.2) | - | - |
| **Hoehn & Yahr stage** | 1.5 (1.0 - 2.0) | - | - |
| **MDS-UPDRS III** | 28.5 (20.5 - 34.5) | - | - |
| **LEDD (mg/day)** | 524.8 (243.4 - 817.4) | - | - |

*Values are presented as median (interquartile range) or mean ± standard deviation. Continuous variables were compared using the Mann-Whitney U test and categorical variables using Fisher’s exact test. Disease characteristics were assessed in the PD group only. BMI data were not available for the control group. BMI: body mass index; IQR: interquartile range; LEDD: levodopa equivalent daily dose; MDS-UPDRS: Movement Disorder Society-Unified Parkinson’s Disease Rating Scale; MMSE: Mini-Mental State Examination; PD: Parkinson’s disease; SD: standard deviation.*

### 2.2. Isolation of total serum EVs

Serum was collected into non-gel tubes, supplemented with protease and phosphatase inhibitors and stored at −80 °C. tEVs were isolated by sequential differential centrifugation followed by two rounds of ultracentrifugation at 120,000 x g for 90 min at 4 °C (Supplementary Methods).

### 2.3. Enrichment of L1CAM+ EVs

L1CAM+ EVs were enriched from tEVs by immunocapture with a biotinylated anti-human L1CAM antibody (CD171, clone 5G3) and streptavidin-agarose resin, followed by elution with glycine-HCl and neutralization with Tris-HCl. The protocol was adapted from Mustapic et al. **(29)** and optimized for compatibility with mass spectrometry (Supplementary Methods).

### 2.4. Characterization of tEVs and L1CAM+ EVs

EVs were characterized by protein quantification (BCA assay), particle concentration (MADLS), TEM, cryo-TEM and immunogold labelling, western blotting for CD63, CD81, TSG101 and L1CAM, and super-resolution imaging (Zeiss Elyra 7, Lattice SIM) of L1CAM and pS129-α-synuclein. Detailed protocols and antibodies are provided in Supplementary Methods.

### 2.5. LC-MS-based lipidomics

Lipids were extracted from EV samples normalized to 100 µg protein using a modified Bligh and Dyer method **(33)**, and analysed by UHPLC coupled to a timsTOF mass spectrometer (Bruker Daltonics). A pooled QC sample was included. Lipids were annotated in MS-DIAL (v5.5) based on accurate mass and MS/MS spectra (Supplementary Methods).

### 2.6. Statistics and bioinformatics

Lipid data were filtered, imputed, normalized and log-transformed before analysis. PD and control groups were compared at the species level by the Mann-Whitney U test with Benjamini-Hochberg correction. PLS-DA models were built in SIMCA 18 and validated by permutation testing and CV-ANOVA. Lipid class, chain-length and functional comparisons, lipid set enrichment and reaction-network analyses were performed in LipidSig 2.0 (Supplementary Methods).

## 3. RESULTS

### 3.1. Isolation and characterization of serum tEVs and L1CAM+ EVs

To determine whether neuronal enrichment refines the lipidomic signal recoverable from blood, we isolated EVs from the serum of patients with PD and healthy controls (n = 20 per group; **Figure 1A**). Total EVs (tEVs) were first obtained by ultracentrifugation. In each group, 10 samples were then analysed as tEVs, and an independent set of 10 was subjected to L1CAM immunocapture to obtain L1CAM+ EVs (N = 40 samples in total). Both fractions were characterized according to MISEV2023 recommendations (**36**) before lipidomic analysis.

**Figure 1.**
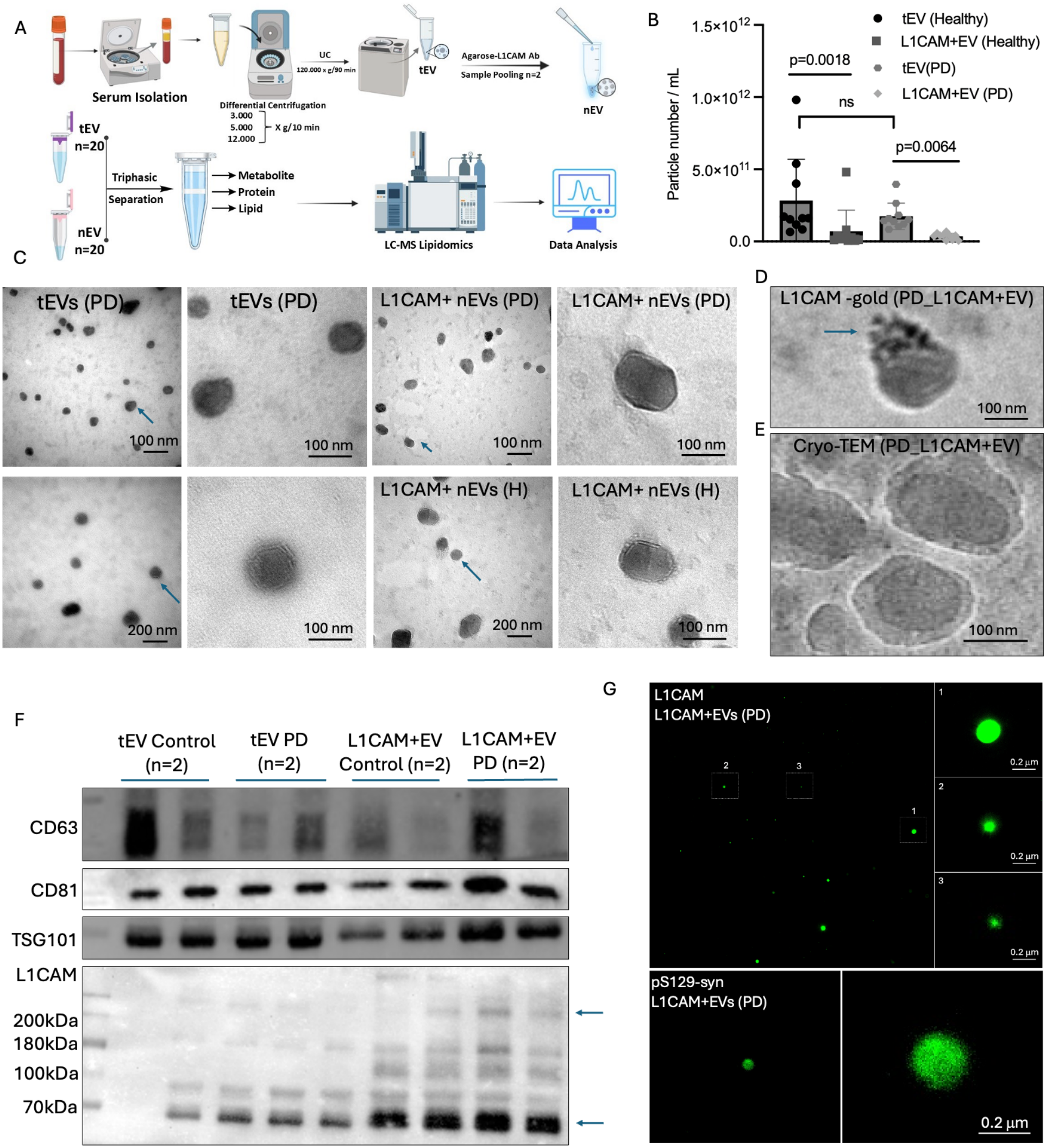
Study design and characterization of serum tEVs and L1CAM+ EVs. **(A)** Schematic overview of the study design. Total EVs (tEVs) were isolated from the serum of healthy controls (HC) and patients with Parkinson’s disease (PD) (n = 20 per group) by ultracentrifugation (UC). In each group, 10 samples were analysed as tEVs and an independent set of 10 was subjected to anti-L1CAM immunocapture (IP) to obtain L1CAM+ EVs. All fractions were analysed by untargeted LC-MS (ESI+) lipidomics (N = 40). **(B)** Particle concentrations of tEVs and L1CAM+ EVs from healthy controls and PD patients, determined by MADLS. Each symbol represents one sample, and bars show mean ± SD. Statistical significance was assessed by one-way ANOVA with Tukey post-hoc; ns, not significant. **(C)** Representative transmission electron microscopy (TEM) images of tEVs and L1CAM+ EVs from PD patients and healthy controls (H). Arrows indicate individual vesicles. **(D)** Immunogold labelling of L1CAM on the surface of an L1CAM+ EV from a PD patient (arrow). **(E)** Cryo-TEM image of L1CAM+ EVs from a PD patient showing intact vesicles with a lipid bilayer. **(F)** Western blot analysis of the EV markers CD63, CD81 and TSG101 and of L1CAM in tEVs and L1CAM+ EVs from controls and PD patients (n = 2 per group). Arrows indicate full-length L1CAM (~200 kDa) and a lower-molecular-weight band (~65-70 kDa). CD63 and CD81 were detected under non-reducing conditions. **(G)** Super-resolution imaging (Zeiss Elyra 7, Lattice SIM) of L1CAM+ EVs from PD patients labelled separately for L1CAM (upper) and pS129-α-synuclein (lower). Numbered insets show magnified individual particles. Scale bars: (C-E) as indicated; (G) 0.2 µm.

MADLS showed particle concentrations of ~4.2 × 10^11^ particles/mL in control tEVs and ~1.4-1.7 × 10^11^ particles/mL in PD tEVs and in the L1CAM+ EV fractions of both groups (**Figure 1B**). The lower particle yield after immunocapture is consistent with the selective recovery of an L1CAM-bearing subpopulation from the total circulating EV pool. Transmission electron microscopy revealed round, membrane-bound vesicles with the characteristic morphology of EVs in both tEV and L1CAM+ EV preparations from PD and control samples. No apparent morphological difference was observed between groups or fractions (**Figure 1C**). Immunogold labelling detected L1CAM on the surface of PD-derived L1CAM+ EVs (**Figure 1D**), and cryo-TEM confirmed intact vesicles with a continuous lipid bilayer (**Figure 1E**).

Western blot analysis showed that the canonical EV markers CD81 and TSG101, and to a variable extent CD63, were present in all fractions (**Figure 1F**). This confirms the vesicular nature of both tEV and L1CAM+ EV preparations. High-molecular-weight L1CAM immunoreactivity (~200 kDa, corresponding to full-length L1CAM) was markedly stronger in L1CAM+ EVs than in tEVs from both PD patients and controls, supporting effective enrichment of L1CAM-bearing vesicles by immunocapture. Super-resolution imaging (Lattice SIM, Elyra 7) further identified individual L1CAM-positive particles with sub-diffraction dimensions in the L1CAM+ EV fraction (**Figure 1G**). Notably, separate labeling with an antibody against α-synuclein phosphorylated at Ser129 (pS129) detected pS129-positive particles in L1CAM+ EVs from PD patients (**Figure 1G**).

### 3.2. PD is associated with an increased lipid-to-protein ratio in circulating EVs, driven predominantly by triacylglycerol species

To compare the overall lipid load of circulating EVs between groups, we first calculated, for each individual, the median log2 abundance of all detected lipid species (n=136) normalized to EV total protein (**Figure 2A,B**). In tEVs, all healthy controls displayed negative median values (approximately −1.2 to −3.7). In contrast, PD samples clustered tightly around zero (approximately −0.5 to +0.7), and the two groups separated completely, with no overlap. Two controls (H9 and H10) showed markedly lower values than the remaining healthy individuals. L1CAM+ EVs showed a shift in the same direction but of smaller magnitude. Healthy controls ranged from approximately −2.4 to −0.2, whereas nine of ten PD samples ranged from approximately −1.0 to +0.5, and the two groups partially overlapped. One PD sample (PD1) exhibited a strongly reduced median lipid level (approximately −6.9).

**Figure 2.**
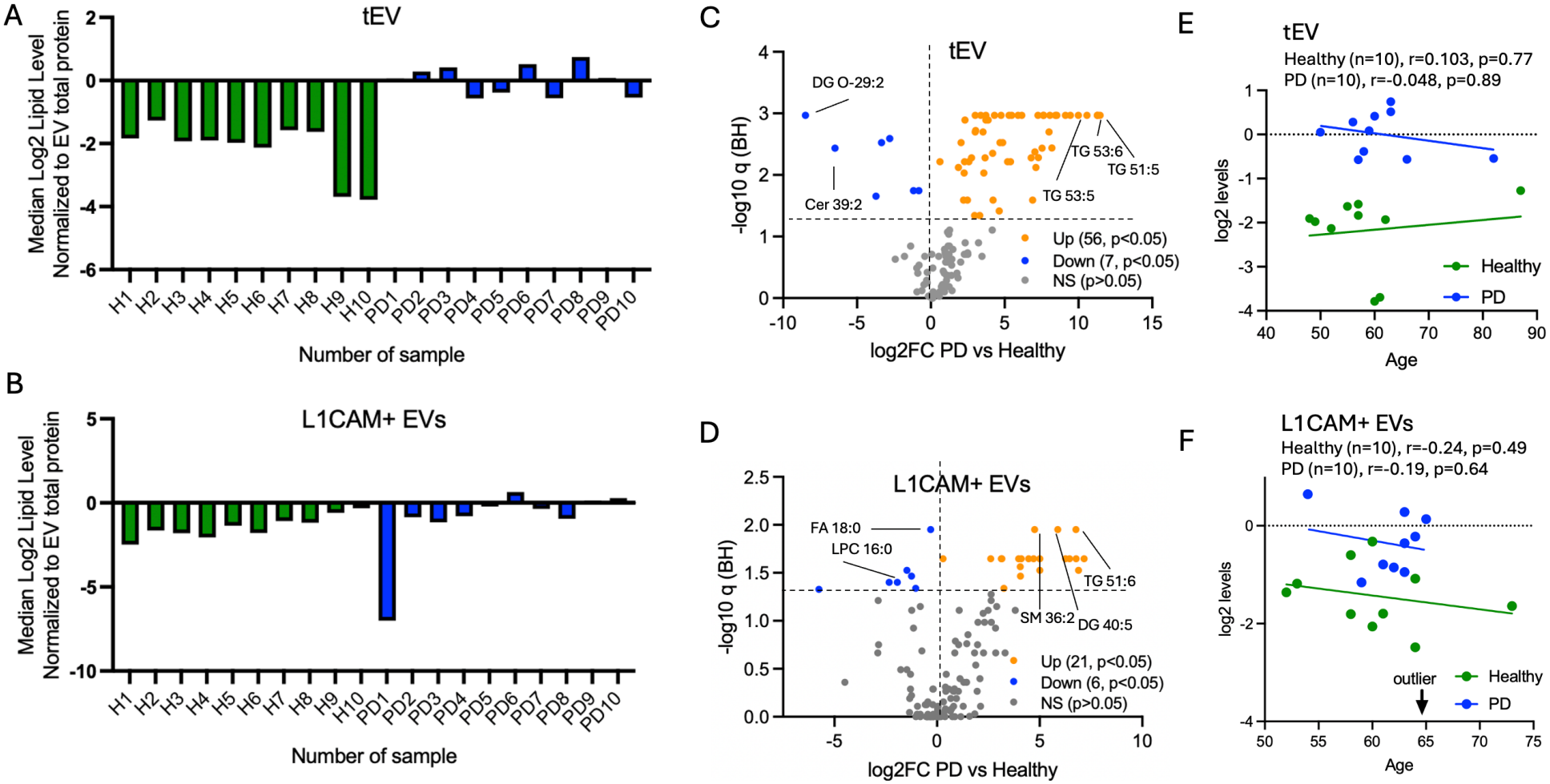
Lipidomic distribution in serum total EVs and L1CAM+ EVs across PD patients versus healthy controls. **(A, B)** Median log2 lipid level of all detected lipid species (n=136) for each individual, normalized to EV total protein, in tEVs (A) and L1CAM+ EVs (B) from healthy controls (H1-H10, green) and PD patients (PD1-PD10, blue). Each bar represents one individual. **(C, D)** Volcano plots of differential lipid abundance between PD patients and healthy controls in tEVs (C) and L1CAM+ EVs (D). The x-axis shows log2 fold change (PD vs healthy), and the y-axis shows −log10 Benjamini-Hochberg-adjusted q value. The horizontal dashed line marks q = 0.05. Orange: significantly increased in PD; blue: significantly decreased in PD; grey: not significant (q ≥ 0.05). Selected species are labelled. **(E, F)** Correlation between donor age and median log2 lipid level in tEVs (E) and L1CAM+ EVs (F), shown separately for healthy controls (green) and PD patients (blue), with linear regression lines. The dotted line marks y = 0. Correlation coefficients (r) and p values were calculated by Spearman correlation. In (F), one PD sample (PD1) falls outside the axis range and is indicated by an arrow “outlier”. n = 10 individuals per group; each of the four groups (tEV/L1CAM+ EV x healthy/PD) comprises samples from independent individuals. TG: triacylglycerol, DG: diacylglycerol, DG O-: ether-linked diacylglycerol, Cer: ceramide, SM: sphingomyelin, FA: fatty acid, LPC: lysophosphatidylcholine.

We next performed differential analysis at the level of individual lipid species (**Figure 2C, D**). In tEVs, 63 species differed significantly between PD and healthy controls (Benjamini–Hochberg-adjusted q < 0.05). Of these, 56 were increased and 7 were decreased in PD. The increased species showed large fold changes (log2FC up to ~11) and were dominated by triacylglycerols, including TG 51:5, TG 53:5 and TG 53:6. The most strongly decreased species were the ether-linked diacylglycerol DG O-29:2 (log2FC: −8.5) and the ceramide Cer 39:2 (log2FC: −6.5). In L1CAM+ EVs, fewer species reached significance (27 in total; 21 increased, 6 decreased), with smaller effect sizes (log2FC up to ~7). Increased species again included triacylglycerols (e.g. TG 51:6), together with SM 36:2 and DG 40:5. Decreased species included FA 18:0 and LPC 16:0.

Because age may influence circulating lipid levels, we tested whether median EV lipid levels correlated with donor age within each group (**Figure 2E,F**). Age ranges overlapped between groups, and we found no significant correlation in either tEVs (healthy: r = 0.103, p = 0.77; PD: r = 0.048, p = 0.89) or L1CAM+ EVs (healthy: r = −0.24, p = 0.49; PD: r = 0.19, p = 0.64). These results indicate that the group differences are not explained by age.

Together, these data show that serum EVs from PD patients carry a higher lipid-to-protein ratio than those from healthy controls. This difference was most pronounced in tEVs, where it was largely attributable to triacylglycerol species, and was smaller in the L1CAM+ EV fraction.

### 3.3. L1CAM-EVs display a distinct lipidomic architecture from tEVs

MADLS showed no significant difference in tEV particle numbers between PD and healthy groups (median 1.47 × 10^11^ vs 1.60 × 10^11^, Kruskal-Wallis followed by Dunn’s test with Bonferroni correction, **Figure 1B**). The lipidomic differences described below therefore reflect changes in vesicle lipid composition rather than in the number of vesicles analysed.

Untargeted lipidomics (**Supplemenary Data I**) followed by PLS-DA clearly separated tEVs from PD patients and healthy controls along the first component (**Figure 3A**). The model showed high goodness of fit and predictive ability (R^2^X = 0.664, R^2^Y = 0.998, Q^2^ = 0.876; Q^2^ intercept: −0.155; CV-ANOVA p-value = 0.000175), indicating a robust, disease-associated tEV lipid signature. The 25 lipid species contributing most to group separation all had variable importance in projection (VIP) scores above 1.2 (**Figure 3B**). The highest-ranking species were free fatty acids (FA 42:10, FA 38:10, FA 20:0, FA 44:10), followed by TG 50:3;O, DLCL 33:2, Cer 36:1;O3, PE 35:5;O, FA 14:0 and PC O-38:4. The discriminating set also included several ceramides (Cer 41:3;O3, Cer 42:2;O3, Cer 43:2;O3), sphingomyelins (SM 41:2;O3, SM 42:3;O3), sphinganine (SPB 18:0;O2), a sulfatide (SL 44:8;O2) and oxidised triacylglycerols. PLS-DA regression coefficients showed that 24 of these 25 species were more abundant in PD tEVs (**Figure 3C**). The largest coefficients belonged to FA 42:10, FA 20:0, TG 50:3;O, FA 38:10 and FA 44:10. TG 53:2;O was the only discriminating species that was lower in PD tEVs.

**Figure 3.**
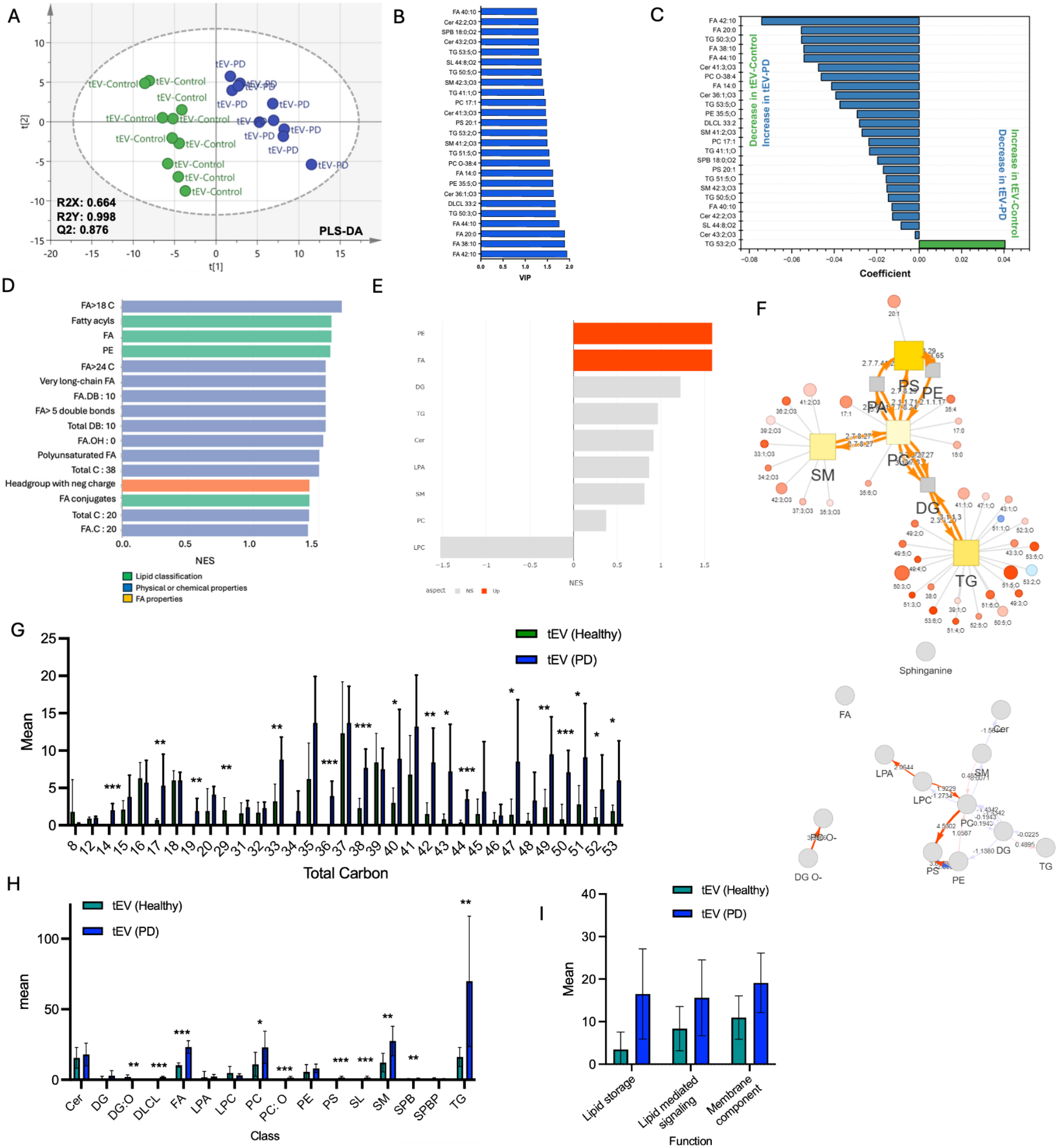
Lipidomic profile of serum tEVs from PD patients compared with healthy controls. **(A)** PLS-DA score plot of tEVs from healthy controls (tEV-Control) and PD patients (tEV-PD) (n = 10 per group). Model parameters (R2X, R2Y and cross-validated Q2) are shown. **(B)** The 25 lipid species with the highest variable importance in projection (VIP) scores. **(C)** PLS-DA regression coefficients of the same species. Negative coefficients indicate higher abundance in PD, positive coefficients higher abundance in controls. **(D)** Lipid ontology enrichment analysis, showing normalized enrichment scores (NES) of significantly enriched terms in PD tEVs, coloured by ontology aspect (lipid classification, physical or chemical properties, fatty acid properties). **(E)** Lipid class enrichment analysis. Classes significantly increased in PD are shown in orange (Up); non-significant classes are shown in grey (NS). **(F)** Reaction-network analysis of PD versus control tEVs. Upper: lipid reaction network showing lipid classes (squares) and individual species (circles), coloured by direction and magnitude of change (red, increased; blue, decreased in PD); edges are labelled with EC numbers of the catalysing enzymes. Lower: pathway activity network showing pathway scores between lipid classes (orange, activated; blue, suppressed in PD). **(G-I)** Abundance of lipids grouped by total carbon number (G), lipid class (H) and lipid function (I) in control and PD tEVs. Data are shown as mean ± SD. Statistical significance was assessed by one-way ANOVA with Tukey post-hoc; *p < 0.05, **p < 0.01, ***p < 0.001.

Lipid set enrichment analysis (LSEA) identified fatty-acid-related terms as the most strongly enriched in PD tEVs (NES ~1.4-1.6; **Figure 3D**). These included fatty acyls with more than 18 and more than 24 carbons, very-long-chain FA, FA with more than five double bonds, polyunsaturated FA, and FA with 10 double bonds. The lipid classes FA and PE, and lipids carrying a negatively charged headgroup, were also enriched. At the class level, FA and PE were significantly enriched in PD tEVs (NES ~1.6; **Figure 3E**). DG, TG, Cer, LPA, SM and PC showed non-significant positive enrichment, whereas LPC showed a non-significant negative trend.

We next compared lipid abundance by total carbon number (**Figure 3G**). PD tEVs contained more lipids at most chain lengths between C33 and C53, which span the phospholipid, sphingolipid and triacylglycerol range. Several species were detected almost exclusively in PD tEVs, including those with C14, C19, C36 and C44 total carbons. Odd-chain lipids (C17, C19, C33, C35, C43, C47, C49, C51, C53) were also increased. C29 was the only chain length that was lower in PD. At the class level (**Figure 3H**), PD tEVs showed increased TG (about 4-fold), SM (about 2.3-fold), FA (about 2.2-fold) and PC (about 2-fold). Low-abundance classes including PS, ether-linked PC (PC O-), dilysocardiolipin (DLCL), sulfatides (SL) and sphingoid bases (SPB) were also increased. Ether-linked DG (DG O-) was the only class decreased in PD tEVs. Cer, DG, LPA, LPC, PE and SPBP abundances did not differ significantly.

Lipid network analysis mapped the changes onto known biosynthetic reactions (**Figure 3F, Supplementary Data II**). The strongest predicted increases in substrate-to-product flux were in the PC-to-PS conversion (phosphatidylserine synthase), LPC-to-LPA conversion (lysophospholipase D/autotaxin) and ether PC-to-ether DG conversion (**Supplemenary Data III**). Reactions linking PS and PE, PC and SM, and DG and TG were also predicted to be upregulated. At the species level, most TG, SM, PC and PS species in the network were increased in PD tEVs.

Grouping lipids by biological function showed that PD tEVs carried more lipids associated with lipid storage (about 4.8-fold), lipid-mediated signalling (about 1.9-fold) and membrane components (about 1.7-fold) than control tEVs (**Figure 3I**).

### 3.4. L1CAM+ EVs display a lipid composition distinct from tEVs in PD

PLS-DA separated L1CAM+ EVs and tEVs from PD patients (R^2^X = 0.733, R^2^Y = 0.999, Q^2^ = 0.925; Q^2^ intercept: −0.135; CV-ANOVA p-value = 1.63 x 10^-5^; **Figure 4A**). The highest VIP scores were obtained for DG O-29:2, FA 20:0, FA 18:0, FA 12:0, SM 41:2;O3, FA 16:4 and SM 41:3;O3 (**Figure 4B**). Most discriminating features were less abundant in L1CAM+ EVs (**Figure 4C**). These included DG O-29:2, saturated FAs (FA 12:0, FA 18:0, FA 20:0), a series of SM species (SM 31:2, 32:2, 33:2, 35:2, 37:2, 39:2, 41:2 and 41:3;O3), PE 37:5 and PE 37:5;O, PC 37:6;O and DLCL 33:2. SPB 18:0;O2, FA 16:4, PC 35:3;O and PC 33:3;O were relatively enriched in L1CAM+ EVs.

**Figure 4.**
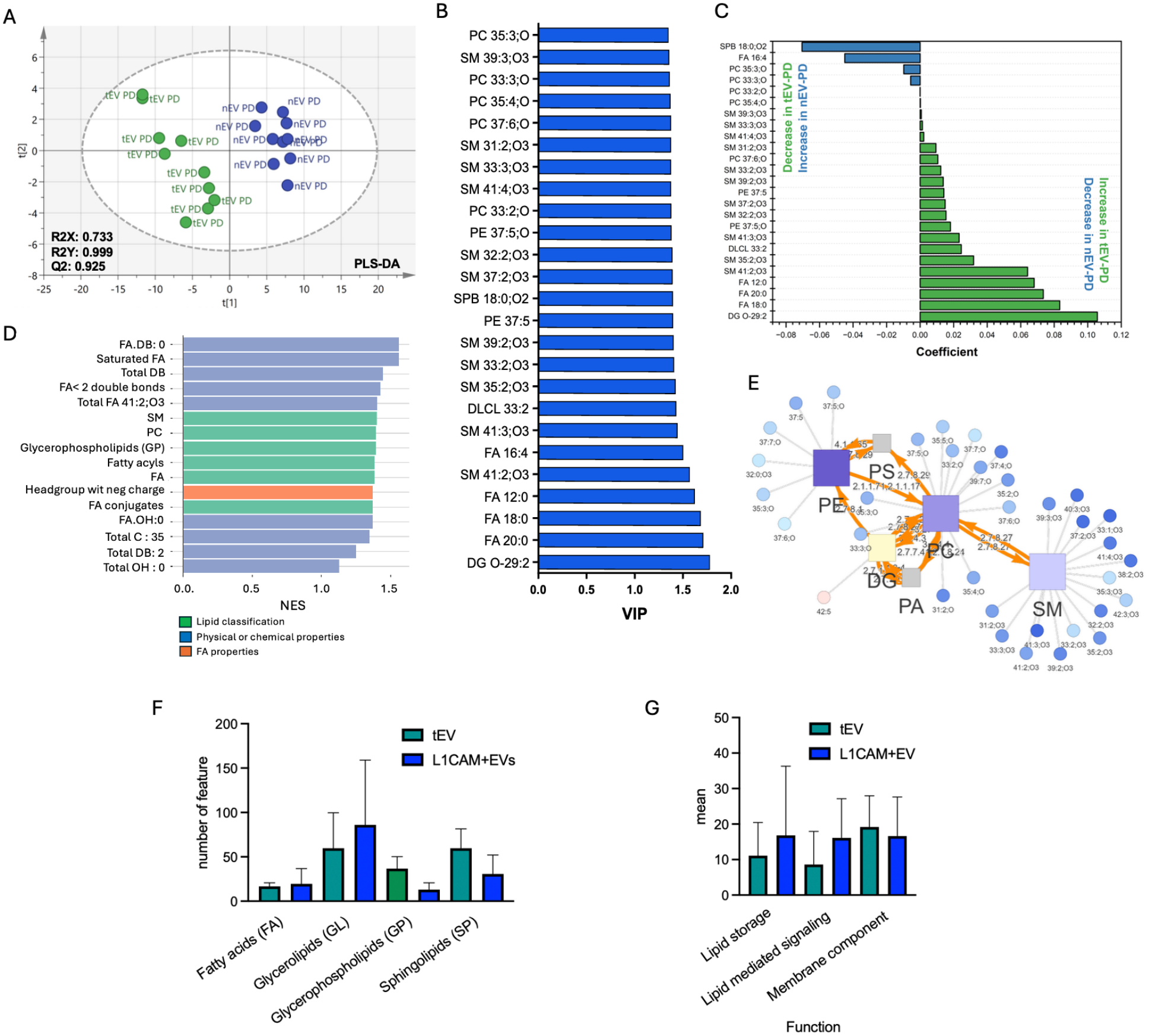
Lipid composition of L1CAM+ EVs compared with tEVs in PD patients. **(A)** PLS-DA score plot of tEVs and L1CAM+ EVs from PD patients (n = 10 per group). Model parameters (R2X, R2Y and cross-validated Q2) are shown. **(B)** The 25 lipid species with the highest VIP scores. **(C)** PLS-DA regression coefficients of the same species. Negative coefficients indicate higher abundance in L1CAM+ EVs, positive coefficients higher abundance in tEVs. **(D)** Lipid ontology enrichment analysis showing NES of terms enriched among lipids lower in L1CAM+ EVs than in tEVs, coloured by ontology aspect (lipid classification, physical or chemical properties, fatty acid properties). **(E)** Lipid reaction network of L1CAM+ EVs versus tEVs. Squares represent lipid classes and circles individual species, coloured by direction and magnitude of change (blue, decreased; red, increased in L1CAM+ EVs). Edges are labelled with the EC numbers of the catalysing enzymes. **(F, G)** Abundance of lipids grouped by lipid category (F) and lipid function (G) in tEVs and L1CAM+ EVs. Data are shown as mean ± SD. Statistical significance was assessed by one-way ANOVA with Tukey post-hoc; *p < 0.05, **p < 0.01.

Enrichment analysis showed that the features lower in L1CAM+ EVs clustered in SM, PC, glycerophospholipids and fatty acyls, in saturated fatty acyl chains (FA.DB:0, FA with < 2 double bonds) and in lipids with negatively charged headgroups (**Figure 4D**). In the reaction network, the depleted species mapped predominantly to the SM, PC and PE nodes and their connecting reactions (**Figure 4E**). At the category level, glycerophospholipids and sphingolipids were lower in L1CAM+ EVs, whereas fatty acids and glycerolipids did not differ significantly (**Figure 4F**). No significant differences were found between functional categories (lipid storage, lipid-mediated signalling, membrane component; **Figure 4G**).

### 3.5. L1CAM+ EVs capture a distinct PD-associated lipid signature

Having established that L1CAM+ EVs differ in lipid composition from tEVs, we next asked whether this fraction captures PD-associated changes and how these compare with those observed in tEVs. PLS-DA clearly separated L1CAM+ EVs from PD patients and controls (R^2^X = 0.711, R^2^Y = 0.998, Q^2^ = 0.879; Q^2^ intercept: −0.0064; CV-ANOVA p-value = 0.0072; **Figure 5A, Supplementary Data IV**), with a predictive ability comparable to that of the tEV model (Q2 = 0.876; **Figure 3A**). The species with the highest VIP scores included saturated fatty acids of odd and even chain length (FA 17:0, FA 18:0, FA 20:0, FA 14:0, FA 15:0), LPA 19:4, FA 40:10, diacylglycerols (DG 40:5, DG 42:5), PS 20:1, odd-chain PC species (PC 15:0, PC 17:0, PC 17:1), several SM species (SM 33:1, 34:2 and 36:2;O3) and SPB 18:0;O2 (**Figure 5B**). Most of these were more abundant in PD L1CAM+ EVs, with the largest coefficients for FA 42:10, SPB 18:0;O2, FA 44:10, FA 17:0 and DG 40:5. In contrast, FA 12:0, FA 18:0, TG 52:3;O, Cer 39:1;O3, LPC 18:2, LPC 16:1 and LPC 16:0 were higher in controls (**Figure 5B, C**).

**Figure 5.**
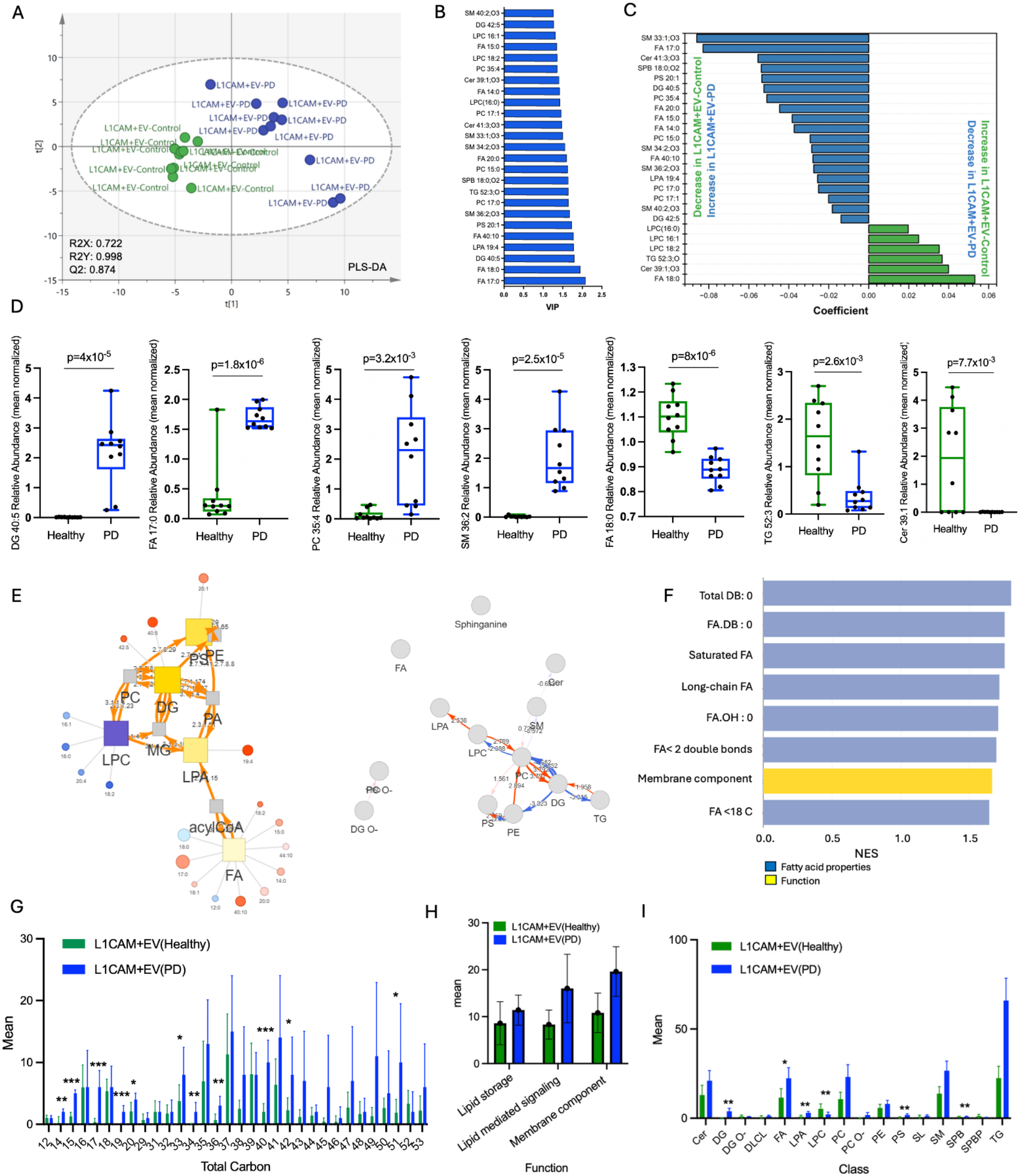
PD-associated lipid alterations in L1CAM+ EVs. **(A)** PLS-DA score plot of L1CAM+ EVs from healthy controls and PD patients (n = 10 per group). Model parameters (R2X, R2Y and cross-validated Q2) are shown. **(B)** The 25 lipid species with the highest variable importance in projection (VIP) scores. **(C)** PLS-DA regression coefficients of the same species. Negative coefficients indicate higher abundance in PD, positive coefficients higher abundance in controls. **(D)** Relative abundance (mean-normalized) of representative discriminating species in L1CAM+ EVs from healthy controls and PD patients. Box plots show the median and interquartile range, whiskers extend from minimum to maximum, and each dot represents one sample. **(E)** Reaction-network analysis of PD versus control L1CAM+ EVs. Left: lipid reaction network showing lipid classes (squares) and individual species (circles), coloured by direction and magnitude of change (red, increased; blue, decreased in PD). Edges are labelled with the EC numbers of the catalysing enzymes. Right: pathway activity network showing pathway scores between lipid classes (orange, activated; blue, suppressed in PD). **(F)** Lipid ontology enrichment analysis showing NES of terms significantly enriched in PD L1CAM+ EVs, coloured by ontology aspect (fatty acid properties, function). **(G-I)** Abundance of lipids grouped by total carbon number (G), lipid function (H) and lipid class (I) in L1CAM+ EVs from healthy controls and PD patients. Data are shown as mean ± SD. Statistical significance in (D) and (G-I) was assessed by unpaired two-tailed Student’s t-test; exact p-values are shown in (D). *p < 0.05, **p < 0.01, ***p < 0.001.

Univariate analysis of representative discriminating species confirmed these differences at the level of individual samples (**Figure 5D**). DG 40:5 (p = 4 x 10^−5^), FA 17:0 (p = 1.8 x 10^−6^), PC 35:4 (p = 3.2 x 10^−3^) and SM 36:2;O3 (p = 2.5 x 10^−5^) were higher in PD L1CAM+ EVs. FA 18:0 (p = 8 x 10^−6^), TG 52:3;O (p = 2.6 x 10^−3^) and Cer 39:1;O3 (p = 7.7 x 10^−3^) were lower. DG 40:5 and SM 36:2;O3 were barely detectable in control L1CAM+ EVs, whereas Cer 39:1;O3 was largely absent in PD samples.

Reaction-network analysis revealed a pattern distinct from that of tEVs (**Figure 5E, Supplementary Data V**). In the lipid reaction network, SM (SM 33:1, 34:2, 36:2 and 40:2;O3) and DG species were increased in PD, whereas LPC species were decreased. Consistent with this, the highest positive pathway scores were obtained for routes that converge on the generation of DG from PC (**Supplementary Data VI**). These included the conversion of PE to PC by PEMT followed by its hydrolysis to DG (4.73), the reacylation of LPC to PC by LPCATs followed by DG formation (4.65), the direct hydrolysis of PC to DG associated with SMPD1 (3.79), and the formation of DG from SM via PC through SGMS1/2 (3.19). The conversion of LPC to LPA by phospholipase D (PLD1-4; 2.54) also scored positively, and PS synthesis by PTDSS1/2 from PE, either directly or via PC, was moderately increased (2.06 and 3.15, respectively). In contrast, pathways that consume DG were predicted to be reduced. These included the de novo synthesis of PC and PE from DG by CEPT1/CHPT1 (−4.65 and −3.32, respectively), the acylation of DG to TG (−3.32), and the synthesis of SM and ceramide downstream of PC (−3.98 and −3.61, respectively). PLA2-mediated hydrolysis of PC to LPC was also predicted to be reduced (−2.39), and this reduction was more pronounced when coupled to de novo PC synthesis from DG (−4.98).

Lipid ontology enrichment showed that the features increased in PD were concentrated in saturated (Total DB: 0, FA.DB: 0, < 2 double bonds), long-chain (≤ 18 carbons) fatty acyl chains without hydroxyl groups (FA.OH: 0), and in lipids annotated as membrane components (**Figure 5F**). Consistent with this, significant differences in total carbon number clustered mainly at C14-C20 and were higher in PD, with additional increases at C33, C34, C36, C40, C42 and C51 (**Figure 5G**). Lipids annotated to lipid-mediated signalling and membrane structure were significantly increased in PD, whereas storage lipids were not (**Figure 5H**). At the class level, PD L1CAM+ EVs contained more DG, FA, LPA and PS and less LPC than control L1CAM+ EVs, whereas TG, SM and PC showed increasing trends that did not reach statistical significance (**Figure 5I**).

## 4. DISCUSSION

In this study, we isolated tEVs and L1CAM+ EVs from the serum of PD patients and age-matched healthy controls and performed untargeted lipidomics to characterise disease-associated changes in the EV lipidome. The results showed distinct lipidomic profiles of tEV and nEVs both separating PD from individuals without parkinsonism.

The characterization data show that both preparations meet the minimal criteria for EV identity (**37**), and that L1CAM immunocapture yields a vesicle population enriched in the neural marker L1CAM. The neuronal specificity of circulating L1CAM-positive vesicles remains debated: a substantial fraction of plasma L1CAM has been reported to be non-vesicular (**34,35**), whereas single-EV analyses support its use as a marker of neuron-derived EVs (**36**). Our observation of membrane-associated L1CAM by immunogold labelling and of L1CAM-positive single particles by super-resolution microscopy indicates that, in our preparations, L1CAM is at least partly vesicle-bound. The detection of pS129-α-synuclein in L1CAM+ EVs from PD patients agrees with previous reports of elevated α-synuclein in neuronally enriched blood EVs in PD and prodromal PD (**28–30**). It suggests that this fraction carries disease-relevant neuronal cargo, and that its lipid composition may therefore reflect neuron-associated changes in PD.

The serum tEVs from PD patients carried a distinct lipid cargo, although their number did not differ from that of healthy controls. The PLS-DA model separated the two groups with high predictive power (Q2 = 0.876), and almost all discriminating species were increased in PD. Because particle numbers were comparable, these findings point to altered lipid loading per vesicle, or a shift in the cellular origin of circulating EVs, rather than increased EV release.

The most prominent feature of the PD tEV lipidome was the enrichment of free fatty acids, particularly very-long-chain polyunsaturated species (FA 38:10, FA 40:10, FA 42:10, FA 44:10). FA longer than C24 are produced by ELOVL4 in a small number of tissues, including the brain and retina (**38**), and are degraded mainly by peroxisomal β-oxidation. Their accumulation in tEVs may therefore reflect increased elongation, reduced peroxisomal clearance, or selective export of these lipids via EVs. This finding is relevant to PD because α-synuclein binds PUFA, and PUFA promotes the formation of soluble aSyn oligomers; these oligomers are elevated in PD and DLB brain (**39**). The enrichment of lipids with negatively charged headgroups is also notable, because aSyn adopts its membrane-bound helical conformation preferentially on anionic membranes (**40**). PD tEVs may thus provide a lipid surface more favourable to aSyn binding and nucleation, in line with evidence that EV and membrane lipids accelerate aSyn aggregation (**41,42**).We also observed increased odd-chain lipids (e.g. C17, C19, PC 17:1). Odd-chain fatty acids derive largely from diet and gut microbial propionate metabolism, and partly from peroxisomal α-oxidation. Given the proposed role of the gut-brain axis in PD, the source of these species warrants further investigation.

TG levels were about 4-fold higher in PD tEVs, and “lipid storage” was the most strongly increased functional category. aSyn binds lipid droplets and promotes TG accumulation (**43**), and aSyn toxicity in yeast and neurons is accompanied by increased unsaturated FA, DG and TG (**44**). Our data are consistent with a systemic shift towards neutral lipid storage in PD. However, serum EV preparations can co-isolate TG-rich lipoproteins (**45,46**). Some of the TG signal may therefore derive from lipoprotein carry-over. Further assessments for measuring ApoB and ApoA1 in the EV fractions are needed.

Another notable finding was that, PD tEVs contained more SM, several hydroxylated ceramides, sulfatides and sphinganine. Sphingolipid dysregulation is a recurring theme in PD: GBA1 variants are the strongest genetic risk factor for the disease, and plasma ceramide levels and ceramide acyl-chain composition are altered in PD patients (**17,47**). Ceramide generated by neutral sphingomyelinase 2 also drives ESCRT-independent ILV formation and exosome release (**48**). The SM increase (about 2.3-fold) was larger than the change in total ceramide, which was not significant. This imbalance raises the possibility that SM-to-ceramide conversion along the EV biogenesis route is altered in PD.

PS, PC and ether-linked PC were increased in PD tEVs. Network analysis also predicted increased PC-to-PS and PS-to-PE flux, consistent with LSEA enrichment of PE. PS is exposed on the EV surface and mediates EV uptake by recipient cells, including microglia. Higher PS content may thus alter how PD EVs are taken up and cleared. Plasmalogen loss has been reported in PD brain lipid rafts (**49**); the opposite trend we observed for ether PC in tEVs may reflect export of these lipids or a different cellular origin, and requires confirmation. The increase in dilysocardiolipin, a cardiolipin degradation product, suggests mitochondrial membrane turnover. This is of interest because cardiolipin exposure on the outer mitochondrial membrane recruits aSyn and modulates its aggregation (**50**).

The lipid composition of L1CAM+ EVs differed markedly from that of tEVs in PD, mainly through lower levels of SM, glycerophospholipids and saturated fatty acids. Two non-exclusive explanations are likely. First, tEVs isolated by ultracentrifugation co-isolate lipoproteins rich in PC, SM and neutral lipids (**25**). The additional affinity-capture and wash steps would reduce this contribution, so part of the difference likely reflects higher purity of the immunocaptured fraction. Second, EV subpopulations of different cellular and biogenetic origin differ in membrane lipid composition, particularly in SM and saturated acyl chains that determine membrane order and rigidity (**20,21,23**). The relative enrichment of SPB 18:0;O2 (sphinganine), an intermediate of de novo sphingolipid synthesis, together with lower SM levels, points to a distinct sphingolipid composition of the L1CAM+ fraction, consistent with a different cellular origin or biogenetic route. Because the two fractions were obtained from independent individuals, inter-individual variation may also contribute. However, the high cross-validated predictive ability of the model indicates that fraction identity is the dominant source of variance. Together, these findings show that L1CAM immunocapture isolates a lipidomically distinct vesicle population rather than a subsample of the tEV lipidome. This is a prerequisite for the possibility that L1CAM+ EVs capture disease-associated changes that are not evident in tEVs.

The central question of this study was whether L1CAM enrichment provides PD-related lipidomic information beyond that available from tEVs. Both fractions discriminated PD from controls with comparable discriminatory ability, indicating that L1CAM enrichment does not improve disease classification per se. However, the PD signature of L1CAM+ EVs differed qualitatively from that of tEVs, so it cannot simply be a diluted or concentrated version of the tEV signal. In L1CAM+ EVs, PD was characterized by accumulation of DG and LPA, depletion of LPC, and a predicted shift of PC metabolism from synthesis and remodelling toward catabolism to DG. Together with the distinct baseline composition of L1CAM+ EVs (**Figure 4**), these results suggest that L1CAM immunocapture enriches for vesicles whose lipid cargo reflects disease-associated processes not evident in the total circulating EV pool.

Several features of the L1CAM+ EV signature are relevant to PD biology. DG and LPA are bioactive lipids involved in signalling, and the selective increase in lipid-mediated signalling categories in PD L1CAM+ EVs suggests that the changes are not merely structural. The network analysis implicated enzymes with established genetic links to PD. SMPD1, encoding acid sphingomyelinase, carries variants associated with PD risk (**44,51**), and its predicted involvement in PC to DG transformation and SM-related routes connects this signature to the lysosomal sphingolipid pathway also highlighted by GBA1 (**15,16**). Likewise, reduced PC to LPC flux, together with lower LPC levels, points to decreased phospholipase A2 activity. This is notable because loss-of-function mutations in PLA2G6 cause early-onset parkinsonism (**52**). In both fractions, the predicted SM to Cer conversion was reduced, and ceramide levels did not increase. This contrasts with the ceramide accumulation described in PD brain (**17**) and in EVs from Lewy body disease (**24**). One possible explanation is that ceramide generated in neurons is not proportionally exported in circulating EVs, or that reduced sphingomyelinase activity limits ceramide-driven vesicle biogenesis (**22**). The shared increase in PS across both fractions suggests that PS enrichment is a common feature of circulating EVs in PD. Given the preferential binding of α-synuclein to anionic phospholipids (**13,14**), this may be relevant to EV-associated α-synuclein propagation.

## 5. LIMITATIONS

This study has several limitations. The cohort was small (n = 10 per group), and the findings have not yet been validated in an independent cohort. The four groups comprised different individuals, so comparisons between L1CAM+ EVs and tEVs were unpaired and may partly reflect inter-individual variation. This comparison was also restricted to PD samples, leaving open whether the compositional differences are intrinsic to the L1CAM+ fraction or shaped by disease. Lipoprotein markers such as ApoA1 and ApoB were not assessed, so the contribution of co-isolated lipoproteins to the lipid profile of tEVs cannot be quantified. The lower lipid input after immunocapture may have pushed low-abundance species below the detection limit. Several class-level differences, particularly for DG and PC O-, arose from lipids near this limit, and presence-versus-absence effects may have amplified their magnitude. Diet, fasting status and dopaminergic medication can influence circulating fatty acids and TG, and were not modelled as covariates. This is particularly relevant to the odd-chain fatty acids, which may originate from diet or gut microbial metabolism. Annotations such as FA 42:10, FA 44:10 and SPB 18:0;O2 are based on sum composition and require confirmation by targeted MS/MS. Finally, pathway scores were inferred from lipid abundance ratios rather than enzyme activity, so the involvement of SMPD1 and PLA2G6 should be regarded hypothetical.

## 6. CONCLUSION

In summary, serum tEVs and L1CAM+ EVs from PD patients carry distinct lipid compositions and distinct PD-associated lipid signatures (**Figure 6**). Both fractions discriminated PD from healthy controls with comparable discriminative ability. In tEVs, PD was associated with increased storage lipids, sphingomyelins and PS synthesis. In L1CAM+ EVs, PD was characterized by accumulation of DG and LPA, depletion of LPC and a predicted shift of PC metabolism toward DG generation. Increased PS and reduced predicted conversion of SM to ceramide were common to both fractions. L1CAM enrichment does not improve disease classification, but it captures qualitatively different, mechanistically interpretable lipid changes. This supports the evaluation of L1CAM+ EV lipidomics as a complementary approach to biomarker discovery in PD, which will require validation in larger, independent cohorts.

**Figure 6.**
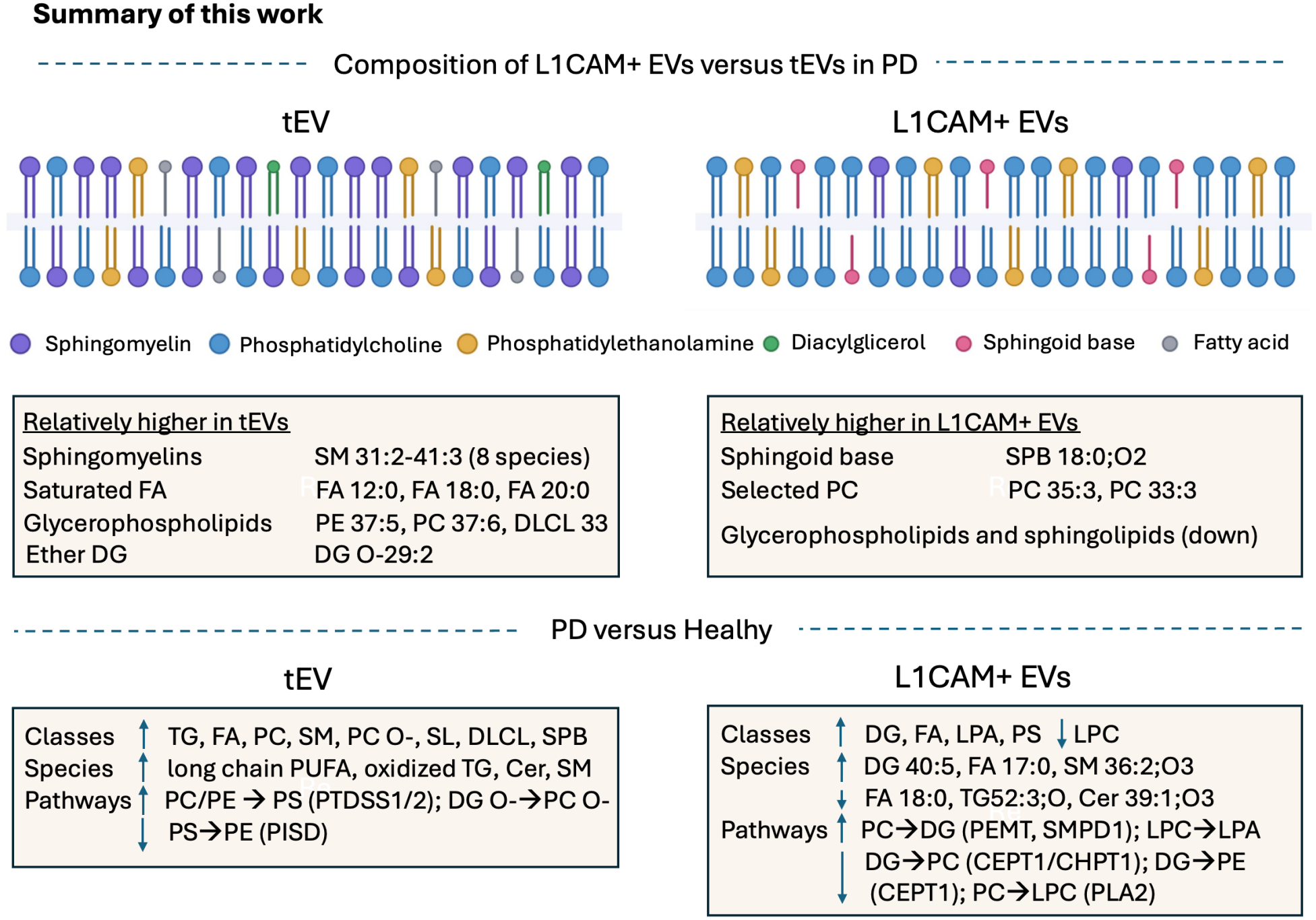
Schematic summary of work. (Upper) Lipid composition of tEVs and L1CAM+ EVs in PD patients. Compared with L1CAM+ EVs, tEVs contained relatively higher levels of sphingomyelins, saturated fatty acids, selected glycerophospholipids and ether-linked DG. L1CAM+ EVs were relatively enriched in the sphingoid base SPB 18:0;O2 and selected PC species, and contained lower levels of glycerophospholipids and sphingolipids overall. Membrane drawings are schematic and highlight only the main compositional differences; leaflet distribution was not measured. (Lower) Lipid alterations in PD compared with healthy controls in tEVs and L1CAM+ EVs, shown at the level of lipid classes, representative species and predicted pathway activity. Up and down arrows indicate increased and decreased levels or pathway activity in PD, respectively.

## Acknowledgement

This study was financially supported by Ankara University Scientific Research Projects Coordination Office (TSA-2025-3825). The mass spectrometry infrastructure used in this study is funded by the Presidency of the Republic of Turkey through the Presidency of Strategy and Budget under grant number 2016 K 121230, and partially by the Hacettepe University Scientific Research Unit (BAP) under project number FAY-2026-22610.

