## Supplementary Methods for "Lipidomic Profiling of L1CAM+ Extracellular Vesicles in Parkinson’s Disease"

### 1.1. Purification of total serum EVs

Peripheral whole blood was collected into serum collection tubes (red-cap tubes; non-gel tubes were preferred for LC-MS-based downstream analyses). Blood samples were allowed to clot at room temperature for 30 min, followed by serum separation according to standard clinical procedures. Immediately after serum collection, protease inhibitor cocktail and phosphatase inhibitor cocktail (Sigma-Aldrich) were added to final concentrations of 1% (v/v) each. Serum aliquots were transferred into low-protein-binding tubes and stored using a stepwise freezing procedure at -20 °C followed by long-term storage at -80 °C until EV isolation.

For tEV isolation, serum samples were thawed on ice and subjected to sequential differential centrifugation at 4 °C to remove cells, cell debris, and large particles: 3000 x g for 10 min, 5,000 x g for 10 min, and 12,000 x g for 10 min. The resulting supernatant was transferred into polypropylene ultracentrifuge tubes (1.5 mL) and ultracentrifuged at 120,000 x g for 90 min using a Himac CP100NX ultracentrifuge (rotor P50A3). The pellet was washed by resuspension in 0.22 µm-filtered phosphate-buffered saline (PBS), followed by a second ultracentrifugation step at 120,000 x g for 90 min under the same conditions. The final EV pellet was resuspended in 100 µL PBS supplemented with protease and phosphatase inhibitors (1% v/v each) and stored at -80 °C until further characterization or downstream analyses.

### 1.2.Enrichment of L1CAM+ extracellular vesicles (L1CAM+ EVs)

For immunocapture, tEVs (200 µL) obtained from 3 mL serum were incubated with 4 µg biotinylated mouse anti-human L1CAM antibody (CD171, clone 5G3; Thermo Fisher Scientific/eBioscience) in a final volume containing 50 µL phosphate-buffered saline (PBS) supplemented with 3% bovine serum albumin (BSA). Samples were mixed thoroughly and incubated at 4 °C overnight under gentle agitation. Subsequently, 15 µL streptavidin-agarose resin (UltraLink; Thermo Fisher Scientific), pre-equilibrated in 50 µL PBS containing 3% BSA, was added to each sample. The suspension was incubated for 4h at 4 °C with continuous mixing to allow capture of antibody-bound vesicles. Resin-bound complexes were collected by centrifugation at 200 × g for 10 min at 4 °C, and the supernatant was carefully removed. Bound vesicles were eluted by adding 200 µL 0.1 M glycine-HCl to the pellet and mixing briefly for 30 s. Samples were then centrifuged at 4,500 × g for 10 min at 4 °C to separate the resin-antibody matrix from the released vesicle fraction. The resulting supernatant, containing enriched L1CAM+ EVs, was immediately transferred to fresh tubes containing 15 µL 1 M Tris-HCl for neutralization and mixed gently. For the mass spectrometry-based downstream applications, BSA was omitted from all buffers whenever possible to minimize protein contamination and ion suppression effects. This sequential workflow, involving UC-based isolation of total EVs followed by L1CAM-targeted EV enrichment, was adapted from the approach described by Mustapic et al. (**29**), while incorporating protocol refinements such as combining tEV inputs, adjusting capture/incubation steps, and replacing standard reagents with mass spectrometry-friendly buffer systems.

### 1.3.Characterization of tEVs and L1CAM+EVs

Total protein concentration is determined using a Thermo BCA kit. For this assay, 5 µL of each isolated EV sample is diluted with PBS to a final volume of 25 µL. Then, 200 µL of the working reagent is added to each diluted sample (final volume 225 µL), followed by 30 min incubation, after which absorbance is measured according to the kit instructions. For DLS measurements, EV samples are diluted by adding 10 µL EV sample to 990 µL of 0.22 µm-filtered PBS (final volume 1000 µL). Size measurements are performed using the MADLS system (Malvern Panalytical, UK**)**.

For microscopic observations, TEM grids (200-mesh, carbon-coated) are used. EV samples are diluted 1:2, and 10 µL of diluted EV sample is deposited onto the grid and incubated for 1 h. Excess liquid is gently wicked off using the edge of a filter paper. Grids are then subjected to a washing step and negative staining using 1% uranyl acetate (0.22 µm-filtered). After staining, grids are imaged without an additional post-stain wash.

WB: EV pellets from tEV and L1CAM+EV fractions were lysed in RIPA buffer containing protease inhibitors, and protein was quantified by BCA assay. Equal amounts of protein (30 µg) were separated on 4-20% Mini-PROTEAN TGX gels (Bio-Rad). CD63 and CD81 were run under non-reducing conditions, and TSG101 and L1CAM under reducing conditions. Proteins were transferred to PVDF membranes using the Trans-Blot Turbo system (Bio-Rad). Membranes were blocked in 5% non-fat milk in TBS-T for 1 h and incubated overnight at 4 °C with primary antibodies against CD63 (Thermo Fisher; 10628D), CD81 (Thermo Fisher; 10630D), TSG101 (ELK Biotech; ES7453) and L1CAM (Biolegend; 371602). They were then incubated with HRP-conjugated secondary antibodies for 1 h. Signals were detected with Clarity Western ECL substrate on a ChemiDoc imaging system and analyzed in Image Lab software (all Bio-Rad).

Imaging of L1CAM+EV using super-resolution microscopy: Isolated L1CAM+ EVs were immobilized on poly-L-lysine-coated high-precision coverslips for 30 min at room temperature and fixed with 4% paraformaldehyde for 15 min. Separate samples were incubated with primary antibodies against L1CAM (Biolegend; 371602) or α-synuclein phosphorylated at Ser129 (pS129) (Abcam; ab209422) overnight at 4 °C. They were then incubated with Alexa Fluor 488/568-conjugated secondary antibodies (1:000) for 1 h at room temperature. L1CAM+ EVs were pelleted by ultracentrifugation at 110,000 × *g* for 70 min at 4 °C, washed once in PBS under the same conditions to remove unbound antibodies, and resuspended in 20 µL PBS. Images were acquired on a Zeiss Elyra 7 system in Lattice SIM mode. Raw data were reconstructed with SIM² processing in ZEN Black, 3.X (Zeiss).

### 1.4.LC-MS-Based lipidomics analyses

*Lipid extraction:* Lipids were extracted from tEV and L1CAM+EV samples (normalized to 100 µg total EV protein) using a slightly modified Bligh and Dyer method (**33**). EV pellets were resuspended in 100 µL ice-cold LC-MS grade water (Sigma-Aldrich). Methanol and chloroform (both LC-MS grade, Sigma-Aldrich) were added sequentially to a final methanol:chloroform:water ratio of 2:1:0.8, v/v/v, and samples were vortexed and incubated on ice for 15 min; phase separation was induced, followed by centrifugation at 5,000 × *g* for 10 min at 4 °C. The lower organic phase was collected, and the pooled organic phases were dried under a vacuum concentrator. Dried extracts were reconstituted in 200 µL isopropanol:acetonitrile:water (6:3:1, v/v/v) and stored at -80 °C until analysis. A pooled quality-control (QC) sample was prepared by combining equal aliquots of all extracts for MS/MS annotation.

*LC-MS analysis*: Lipid extracts were analyzed on a UHPLC system (ELUTE PLUS UHPLC, Bruker Daltonics, Germany) coupled to a mass spectrometer (TIMS-TOF-MS, Bruker Daltonics, Bremen, Germany). Separation was performed on a C18 Intensity Solo 2 column (2.0 µm, 2.1 x 100mm, Bruker Daltonics, Germany) at 50 °C, using mobile phase A (acetonitrile:water (60:40, v/v)) and mobile phase B (isopropanol:acetonitrile (90:10, v/v)), both containing 10 mM ammonium formate and 0.1% formic acid. The gradient program was performed 0-1 min at 35% B; 1-7 min at 55% B; 7-10 min at 75% B; 10-18 min at 95% B; 18-25 min at 35% B at a flow rate of 0.4 mL/min, and the injection volume was 20 µL. The ESI parameters were set as follows: capillary voltage 3000 V, nebulizer pressure 2.2 bar, drying gas flow 10 L/min, and drying gas temperature 220 °C. Mass spectra were acquired in negative and positive ion modes across an *m/z* range of 100-1700 at 2.0 Hz. MS/MS experiments were carried out with precursor selection at 6-12 Hz, a cycle time of 0.5 s, an isolation width of 4.0, and a fixed collision energy of 10 eV, 20 eV, and 40 eV to support lipid annotation. MS/MS data were acquired over *m/z* 50-1700. Raw data were processed in MS-DIAL (ver. 5.5) for peak picking, alignment, and lipid annotation based on accurate mass and MS/MS fragmentation.

### 1.5. Statistics and bioinformatics

Lipid features detected in fewer than 50% of samples in a group were removed, and the remaining missing values were imputed with one-fifth of the minimum value of each feature. Intensities were normalized to internal standards and $\log_{10}$-transformed.

Lipidomic analyses were performed separately for tEV (10 healthy controls, 10 PD patients) and L1CAM+ EVs (10 healthy controls, 10 PD patients). For each EV type, the intensity of each of the 136 annotated lipid species was scaled to the mean intensity of that lipid across all samples, so that a value of 1 corresponds to the cohort mean. Normalized values were then log2-transformed (0 = cohort mean; +1 = twofold above; -1 = twofold below). To summarize the overall lipid content of each sample, we calculated a per-sample lipid level, defined as the median of the log2-normalized abundances across all 136 lipid species (log2 level). For each lipid species, PD patients were compared with healthy controls using the two-tailed Mann-Whitney U test. Fold change was expressed as log2FC (PD − healthy), calculated as the difference between the group means of the log2-normalized abundances. P values were corrected for multiple testing across the 136 lipid species within each EV type using the Benjamini–Hochberg false discovery rate (FDR) procedure. Lipids with $q < 0.05$ were considered significantly altered and classified as increased (log2FC > 0) or decreased (log2FC < 0). The relationship between age and the lipid level was assessed separately in healthy controls and PD patients using Spearman's rank correlation (two-tailed, exact P values). Age distributions between healthy controls and PD patients were compared with the two-tailed Mann-Whitney U test.

Multivariate analysis was performed in SIMCA, version 18 (Sartorius). Partial least squares-discriminant analysis (PLS-DA) was performed for three comparisons: PD versus control tEVs, PD versus control L1CAM+ EVs, and L1CAM+ EVs versus tEVs in PD. Model quality was assessed by $R^2X$, $R^2Y$, and $Q^{2,}$ and model validity and reliability were further evaluated by permutation testing (100 permutations) and cross-validated predictive residuals (CV-ANOVA). Variables with a variable importance in projection (VIP) score > 1.0 were considered discriminating, and the direction of change was determined from the PLS-DA regression coefficients. Lipid characteristics were analyzed using LipidSig 2.0. An adjusted p-value < 0.05 was used as the significance threshold for all analyses. Group differences in lipid abundance were assessed by lipid class, total carbon number, and functional category using t-tests. Lipid set enrichment analysis was used to identify enriched ontology terms. Reaction networks and pathways were analyzed using the LipidSig network module.

Nanoparticle count and lipid class, chain-length and functional summaries were plotted and analysed in GraphPad Prism (version 11). Group differences in particle concentration were assessed by one-way ANOVA with Tukey's post hoc test. Data are presented as mean ± SD. A p-value $< 0.05$ was considered statistically significant (*$p < 0.05$, **$p < 0.01$, ***$p < 0.001$).
